# Glucocerebrosidase rescues alpha-synuclein from amyloid formation

**DOI:** 10.1101/363986

**Authors:** M.S. Barber, H.M. Muller, R.G. Gilbert, A.J. Baldwin

## Abstract

Aggregation of the protein *α*-synuclein (*α*Syn) Into amyloid fibrils is associated with Parkinson’s disease (PD), a process accelerated by lipids. Recently, the lysosomal protein glucocerebrosidase (GCase) has been identified as a major risk factor in both genetic and sporadic PD. Here, we use solution state NMR to reveal that GCase directly inhibits lipid induced *α*Syn amyloidogenesis. Structurally, we show that the mechanism for this requires competition between lipids and GCase for *α*Syn, binding the N and C termini respectively. The affinity of GCase for the C-terminus of *α*Syn is such that not only does it inhibit lipid induced amyloid formation, but also it destabilizes mature *α*Syn amyloid fibrils. These results reveal a competitive molecular “tug-of-war” for *α*Syn termini by GCase and lipid, providing a mechanistic link between the clinically observed links between changes in GCase abundance and Parkinsons disease.

## INTRODUCTION

Parkinson’s disease (PD) and other synucleinopathies are a group of neurodegenerative disorders characterized by the aggregation of *α*-synuclein (*α*Syn) into amyloid rich bodies known as Lewy Bodies (1). Recent clinical studies have linked PD to the lysosomal lipid lytic enzyme Glucocerebrosidase (GCase). Mutation in GCase is the leading genetic risk factor for PD, with 5-10 % of PD patients carrying one of hundreds of possible GCase mutations (2)(Fig 1A). In sporadic PD, wild type GCase is co-localised with Lewy bodies and depleted in the neuronal cells of the *substantia nigra*, the brain structure associated with PD(3–7). Overall, GCase mutation increases the risk of PD by up to 30 fold (8, 9).

**Figure 1:**
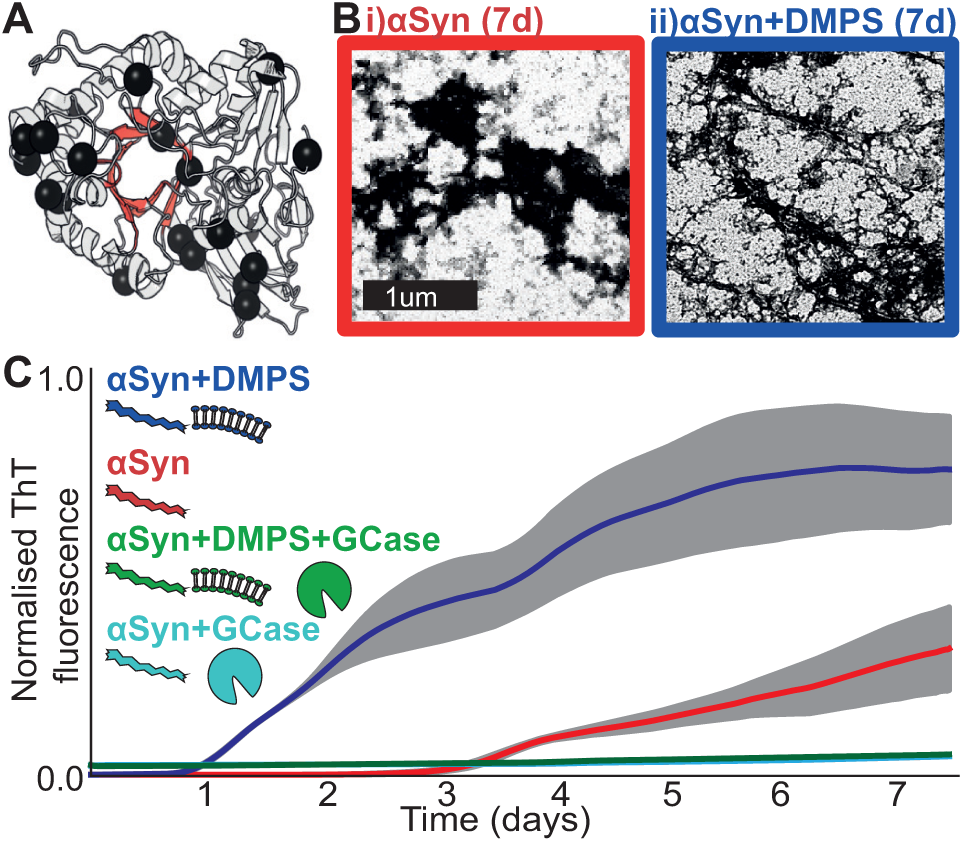
**A**) A structure of GCase (PDB 1OGS) showing mutation sites linked to PD (60) (black), together with the active site (red). **B**) Negatively stained electron micrographs reveal aggregation of *α*Syn and *α*Syn+DMPS samples at pH 5 after 7 days. Though they stain with ThT and the aggregates have a fibrillar aspect, discrete fibrils were not observed. **C**) *α*Syn aggregation kinetics were monitored by a thioflavin-T (ThT) assay at pH 5 with DMPS (blue), GCase (cyan) and both (green). Aggregation of *α*Syn (red) is promoted by DMPS (blue) and inhibited by GCase both with and without DMPS (cyan/green). The grey band shows the standard deviation of 3 independent measurements.

Cellular and animal studies further link GCase to PD. Animal studies of PD demonstrate that knock down of wild type GCase or over expression of mutant GCase causes accumulation of *α*Syn. Likewise, over expression of wild type GCase can reverse *α*Syn related pathological and behavioural features in mouse models (10–13). Healthy cells can degrade both *α*Syn monomers and oligomers through autophagic pathways which culminate in the lysosome (6, 14–17). Lysosomal GCase is depleted in both sporadic and non-sporadic PD (4, 5, 5, 18–24) and intercellular transfer of potentially toxic *α*Syn oligomers between lysosomes is enhanced by GCase depletion (20, 25, 26). Finally, *α*Syn has been shown to interact with GCase directly at lysosomal pH to inhibit GCase activity (27, 28). Taken together, these results suggest a molecular relationship between GCase and *α*Syn that is pertinent to PD. Here we seek to investigate the mechanism by which GCase can rescue *α*Syn from amyloid formation.

At a molecular level, GCase is a 60 kDa glycoprotein found in the acidic lysosome, where it metabolises glucocerebroside (29). Interestingly, post-translational modifications are vital for GCase activity and stability, and ablation of its four N-linked glycosylation sites leads to an inactive and unstable enzyme (30). Therefore, to understand the interplay between GCase and *α*Syn, it is necessary to work with human cell derived GCase complete with relevant post-translational modifications (see methods).

The intrinsically disordered *α*Syn plays a role in synaptic vesicle fusion (31) and forms *α*-helical arrays on the surface of small unilamellar vesicles (SUVs) (32, 33). The sequence of *α*Syn can be divided into three distinct regions, an N-terminal lipid binding domain, a central pro-amyloidogenic domain and a C-terminal domain that remains free in the lipid bound form(33, 34). *α*Syn aggregates into amyloid fibrils that are characterized by ThT binding, increased *β*-strand content, and cross-*β* fibre diffraction patterns (35–38). Notably, this aggregation is accelerated 1000 fold in the presence of membrane mimicking anionic lipids such as DMPS and POPG (32, 39–44), amyloidogenesis is further enhanced in low pH environments (like those found in the lysosome) (45–47). The link between *α*Syn, GCase, acidity and lipids suggest the lysosome as a likely organelle for the onset of *α*Syn pathogenesis (6, 11, 14, 15, 17, 20, 23, 26).

To understand how GCase might influence amyloidosis, we here investigate the relationship between lipids, human cell derived, wild-type GCase and full length *α*Syn under lysosomal conditions by means of solution state nuclear magnetic resonance (NMR) spectroscopy. This technique allows examination of multiple binding partners with transient interactions at atomic resolution. In its protonated form, GCase’s relatively high molecular weight would be expected to give unobservably broad resonances. NMR labelling techniques designed to overcome this barrier(48) rely on prokaryotic expression systems which in this case would not provide protein with the required glycan modifications neccessary for study of a physiologically relevant interaction. Building on recent advances(49), we have developed a method for isotope incorporation and assignment of high molecular weight proteins allowing us to characterise human cell-line derived GCase by NMR. ^1^H-^15^N HSQC (*α*Syn) and ^1^H-^13^C HMQC (GCase) spectra were recorded over a range of concentrations in the presence and absence of detergent micelles formed from SDS, tween, taurochlorate and physiologically relevant SUVs formed from POPG and DMPS (Fig S1). Phosphatidylserines such as DMPS are found in lysosomal inner leaflets and synaptic vesicles (50, 51). These lipids and detergents cover a range of charges and chain lengths and have been previously used to study *α*Syn(39–43, 50, 51). By carefully quantifying the data and relating it to binding models with rigorous care to avoid overfitting, we determine the precise mechanism by which lipds, GCase and *α*Syn interact.

Our NMR results reveal that under lysosomal conditions, GCase and anionic lipids compete for opposite termini of *α*Syn. The differential affinities of GCase and anionic lipids for the N- and C-termini allow GCase to rescue *α*Syn from both lipid binding and subsequent amyloid formation. We show that under these conditions, GCase can destabilise mature *α*Syn fibrils and that *α*Syn contacts GCase near its active site, providing a structural rationale for GCase inhibition by *α*Syn. Taken together, these results reveal that under lysosomal conditions, GCase is a potent chaperone, able to inhibit aggregation of *α*Syn, providing a molecular explanation for its connection to PD.

## MATERIALS AND METHODS

### Expression and purification of *α*-synuclein

Human sequence *α*Syn plasmid was kindly provided by Ad Bax (NIH)(52). Recombinant ^15^N enriched *α*Syn was expressed from BL21(DE3) star cells (ThermoFisher, Waltham, Mass.) containing kanamycin resistant, T7, IPTG inducible plasmids (pET 41, Novagen, Madison, Wisc.). Cells were grown in M9 minimal media; consisting of 6 g/L Na_2_HPO_4_.7H_2_O, 3 g/L KH_2_PO_2_, 0.5 g/L NaCl, 1 mM MgSO_4_, 300 *μ*M CaCl_2_ supplemented with 0.5g/L ^15^NH_4_Cl. Cells were induced at OD_600_0.7 with 1mM IPTG (Sigma-Aldrich, St. Louis, Miss.) for 3 hours at 37°C. After induction cells were pelleted by centrifugation (Avanti J26XP, Beckman Coulter, Brea, Cali.) and stored at −80°C.

Cell pellets were suspended in 50 mM Tris-HCl, pH7.4, 0.5 M NaCl containing protease inhibitors (EDTA free protease inhibitor cocktail, Roche, Basel, Switzerland) and heated to 90°C for 10 minutes in a water bath (VWB2, VWR, Radnor, Penn.). Resulting lysate was clarified by centrifugation at 20,000g for 30 minutes at 4°C and diluted into buffer A (50mM Tris-HCl, pH 7.4). Diluted lysate was loaded onto a 5 mL Hitrap Q column (Sigma-Aldrich, St. Louis, Miss.) pre-equilibrated with buffer A. Protein was eluted with a gradient of buffer B (50mM Tris-HCl, pH 7.4, 1 M NaCl). Fractions with a high absorbance at 280 nm were assayed by SDS-gel electrophoresis, pooled, concentrated and run on a Superdex S75 size exclusion column (Sigma-Aldrich, St. Louis, Miss.). Elution of *α*Syn occurred at a position equivalent to a 60 kDa folded protein, which is consistent with a disordered protein. Supporting this observation is the diffusion coefficient and chemical shifts observed by NMR, both of which are consistent with monomeric disordered protein. Subsequent purity was assayed by SDS-gel electrophoresis. Pure fractions were pooled, buffer exchanged and concentrated into 50 mM citric acid, 100 mM sodium phosphate, 2 mM sodium azide, pH 5 with Amicon concentrators (Millipore, Burlington, Mass.), and stored at −80°C for future use.

### Source of GCase

GCase (VPRIV) was a gift from Shire Pharmaceuticals PLC (Dublin, Ireland). VPRIV is an enzyme replacement therapeutic for Gaucher’s disease. VPRIV is produced from human cell lines supplemented with kifunensine resulting in high mannose glycans favourable for enzyme replacement therapy(53).

### Lipid and detergent preparation

Single unilaminar vesicles (SUVs) and micelles were produced as previously outlined (44). It is important to have a preparation that does not precipitate and is kinetically stable under the solution conditions investigated. Stock solutions of DMPS, POPG, SDS and taurocholate were prepared in 50 mM citric acid, 100 mM sodium phosphate, 2 mM sodium azide (pH 5). DDM was mixed in with POPG and SDS to reduce the affinity of *α*Syn providing more signal intensity in the spectrum. Stock solutions were mixed at 50°C in a Thermomixer (ThermoFischer, Waltham, Mass.) for 4 hours, freeze thawed with a combination of a 50°C water bath (VWB2, VWR) and dry ice 5 times, before sonication for 5 minutes. Stock solutions were then immediately diluted to their working concentration. All lipid concentrations are stated where they were used in the manuscript, and all are above the critical concentration of the specified lipid/detergent.

### ^13^CH_3_ methyl labelling of GCase

It was necessary for this work to devise a labelling protocol for human cell line derived GCase. Labelling of arginine and lysine residues was accomplished by incubating with ^13^C formaldehyde, using methods previously described(54). GCase was buffered in 50 mM citric acid, 100 mM sodium phosphate, 2 mM sodium azide (pH 5) or 50 mM citric acid, 100 mM sodium phosphate, 2 mM sodium azide (pH 7). Borane dimethylamine complex (Sigma-Aldrich, St. Louis, Miss.) was added to dilute (10 *μ*M) GCase at 4°C to a final concentration of 20 *μ*M. Immediately, ^13^C formaldehyde (Sigma-Aldrich, St. Louis, Miss.) was added to a final concentration of 40 *μ*M and the solution incubated on ice at 4°C for 2 hours. This was repeated once and incubated for a further two hours. Finally, Borane dimethylamine is added to a final concentration of 10*μ*M and the reaction incubated overnight at 4°CC. The reaction was quenched by the addition of Trizma (pH 7) (Trizma, Sigma Aldrich, St. Louis, Miss.) to a final concentration of 100 mM. Reaction byproducts are then removed using Amicon concentrators (Millipore, Burlington, Mass.), washing with a buffer containing fresh 50 mM citric acid, 100 mM sodium phosphate, 2 mM sodium azide (pH 5) buffer.

### LCMSMS and Methyl assignment

In order to assign the observed residues, LCMSMS experiments were performed. Peptides were re-suspended in 5% formic acid and 5% DMSO. They were separated on an Ultimate 3000 UHPLC system (ThermoFischer, Waltham, Mass.) and electrosprayed directly into an QExactive mass spectrometer (ThermoFischer, Waltham, Mass.) through an EASY-Spray nano-electrospray ion source (Thermofischer, Waltham, Mass.). The peptides were trapped on a C18 PepMap100 pre-column (300 *μ*m i.d. x 5 mm, 100, ThermoFisher, Waltham, Mass.) using solvent A (0.1% Formic Acid in water) at a pressure of 500 bar. The peptides were separated on an in-house packed analytical column (75 *μ*m i.d. packed with ReproSil-Pur 120 C18-AQ, 1.9 *μ*m, 120, Dr. Maisch GmbH) using a linear gradient (length: 60 minutes, 7% to 28% solvent B (0.1% formic acid in acetonitrile), flow rate: 250 nL/min). The raw data was acquired on the mass spectrometer in a data-dependent mode (DDA). Full scan MS spectra were acquired in the Orbitrap (scan range 350-2000 m/z, resolution 70000, AGC target 3e6, maximum injection time 100 ms). After the MS scans, the 20 most intense peaks were selected for HCD fragmentation at 30% of normalised collision energy. HCD spectra were also acquired in the Orbitrap (resolution 17500, AGC target 5e4, maximum injection time 120 ms) with first fixed mass at 180 m/z.

NMR spectra acquired at pH 5 of GCase (VPRIV) pre-treated with formaldehyde at either pH 5 or pH7 under otherwise identical conditions yielded different spectra (Fig S5). In both cases, only one methyl addition was identified by LCMSMS to present under each condition as described in the text. This led to unambiguous assignment of R120, the only isolated, well dispersed peak structurally proximal to the active site E340 and E235 residues (55). K346, the other methyl identified by LCMSMS led to no obvious change in the NMR spectrum, suggesting its resonance overlays significantly in the centre of the spectrum.

### GCase activity assays

GCase activity was performed in triplicate in 96-well clear flat bottom black polystyrene microplates (Product #3904, Corning) with 120 *μ*M substrate (resorufin-*β*-D-glucopyranoside (Sigma-Aldrich, St. Louis, Miss.)) and 50 nM GCase buffered in 176 mM K_2_HPO_4_, 10 mM sodium taurocholate, 50 mM citrate, 0.01% tween 20, pH 5.9. The final well volume was 0.1ml. Fluorescence of hydrolysed substrate was measured on a Flurostar Omega (584/630 nm excitation and emission respectively) (BMG Labtech, Ortenberg, Germany) at 37°C (56).

### Aggregation assays

Protein aggregation was observed via thioflavin-T (ThT) fluorescence in 96 microwell plates and a Fluostar Optima plate reader (BMG Labtech, Ortenberg, Germany). Microwell plates containing *α*Syn were aggregated either at 200 *μ*M, 45°C or 100 *μ*M, 30°C pH 5 in 50 mM citric acid, 100 mM sodium phosphate, 2 mM sodium azide pH 5. An *in*-*situ* ThT binding assay with 10 *μ*M ThT and 440/490 nm excitation/emission filters monitored aggregation. ThT assays were repeated in triplicate yielding standard deviations for error measurements (Fig 1).

### NMR experiments

NMR experiments were performed on a Varian 600 MHz spectrometer with a 5-mm z-axis gradient triple resonance room temperature probe at 30°C. Isotopically labeled ^15^N *α*Syn and ^13^C GCase were buffered in 50 mM citric acid, 100 *μ*M sodium phosphate, 2 mM sodium azide (pH 5) and concentrated to 100 *μ*M using Amicon concentrators (Millipore, Burlington, Mass.). 2D ^15^N-^1^H HSQC spectra were recorded with 128 scans per transient with acquisition times of 67.5 ms, over 100 complex points and a 1 s relaxation delay. 2D ^13^C-^1^H HMQC were recorded with 64 scans per transient and an acquisition time of 0.064 s, over 100 complex points and a 1.5 s relaxation delay.

Diffusion pulsed field gradient experiments were used to characterize diffusion behaviour. The translational diffusion of specific resonances was determined using ^1^H pulsed field gradient NMR experiments for all mixtures of *α*Syn, GCase and lipids/detergents described in the paper. In such experiments, signal intensity is attenuated according to the strength of an applied gradient (expressed here with respect to the maximum applied gradient strength) in a manner defined by the translational diffusion coefficient. Increased signal attenuation reveals a larger diffusion coefficient. PFG experiments were carried out over 11 gradient strengths with 256 scans per transient and an acquisition time of 0.5 s. The maximum gradient strength was 60 G cm^−1^. The gradient duration was 0.002 s with a 0.15 s delay between gradients. NMRPipe was used to analyse spectra (57) and Sparky was used to pick peaks and prepare figures. Peaks were assigned by comparison to previously published assignments of *α*Syn (58). As not all peaks were assigned due to overlap, we have in total a 60% assignment of resonances that are clearly isolated, whose coverage that uniformly spans the entire sequence. Peak intensity was calculated from peak heights obtained using Sparky. C-terminal and N-terminal intensities represent the mean and standard deviation of the 20 most C or N terminal residues.

Isotope (^15^N) enriched *α*Syn fibrils were pooled and diluted in 50 mM citric acid, 100 mM sodium phosphate, 2 mM sodium azide (pH 5) to a final 100 *μ*M concentration. Stock samples were then split into two equal working samples. Labelled (^13^CH_3_) GCase was added to a single sample to a final concentration of 100 *μ*M. 2D ^1^H-^15^N HSQC and ^1^H-^13^C HMQC data were collected as described above.

### Electron microscopy

Carbon coated grids (Electron Microscopy Sciences, La Sagne, Switzerland) were incubated with diluted microwell sample (10 *μ*M) for 2 minutes and post stained with 2% aqueous uranyl acetate (10 s) and washed with water droplets. Grids were then imaged on a transmission electron microscope (FEI Tecnai 12) operated at 120kV using a digital CCD camera (Ultrascan US1000, Gatan).

### Thermodynamic modelling

The reduction in signal intensity was recorded for the first and last 20 residues of *α*Syn, owing to their highly similar reduced relative intensities, together with GCase as a function of changes in concentration of the three components in the system, *α*Syn, GCase and lipids. The changes in signal intensity were concentration dependent and so we sought an equilibrium-binding model that accounted for all data. The total concentrations were varied of each such that we had 7 measurements of both GCase, N and C-terminal *α*Syn signal, providing 21 intensity datapoints for each of the anionic lipids and detergents DMPS, SDS/DDM and POPG/DDM. The specific concentrations used were such that this number of datapoints is more than sufficient to characterise the specific *K_D_*s.

There are two challenges to account for these observations. The first is to identify which species are forming, and the second is to account for how the formation of these species affects the NMR signal intensity. We sought to determine the model that gives the best fit to the data, subject to the restraint that improved fitting from more complex models is tested for statistical significance by applying an F-test and determining the probability that this improvement came by chance. A <5% probability level was taken as the condition for statistical significance.

The problem involves three species, lipid SUVs/micelles, *α*Syn and GCase. From the three components, there are 5 binary combinations that can be formed. Should we choose to distinguish independent sites for binding on the N and C termini, the formation of each of these can be characterized by the following association equilibrium constants and definitions:

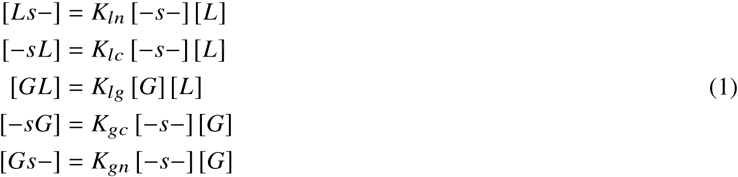

Where the representation -S- for *α*Syn denotes either an empty N/C termini. Replacing one of the ‘−’ symbols with either an L (lipid) or G (GCase) implies occupancy. There are 4 corresponding ternary complexes, which can be similarly defined in terms relevant equilibrium constants:

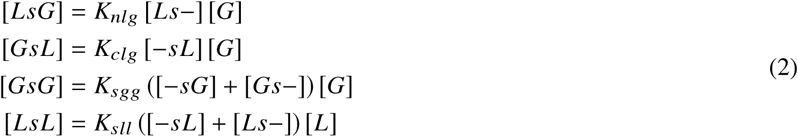

The equilibria are restrained by total numbers of molecules through the following three mass balancing equations:

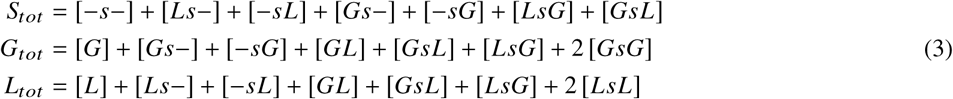

Where S_*tot*_, G_*tot*_ and L_*tot*_ are the total concentrations of *α*Syn, GCase and lipids respectively. For known values of the total concentrations, and trial values of the relevant equilibrium constants, we can self-consistently solve these equations to obtain the corresponding mole fractions of the individual species. By optimising the equilibrium constants against experimental measures of mole fractions, we can obtain their most probable values.

To do this, it is necessary to obtain expressions for the observed signal intensity. There are a number of ways in which this can be formulated depending on the precise details of timescales of the interactions. The timescale together with the specific structural changes that accompany the association will affect the signal intensities through chemical exchange. As described in the text, as we observe neither appreciable line broadening nor change in the chemical shift of resonances, we expect slow or slow-intermediate exchange. Moreover, no variation in the R^2^ of the residues was observed with 180° pulse frequency in CPMG experiments, suggesting no species interconversion on the milli-second exchange.

If a species rigidly tumbles like either GCase or SUV/micelle, we would not expect it to contribute directly to the solution NMR spectra. Nevertheless, if part of *α*Syn retains some of its conformational flexibility when bound, we would expect it to contribute to the spectrum, although with a reduced signal intensity. To account for this type of interaction, and to first order to account for the effects of chemical exchange, the relaxation correction factors *γ* are introduced such that we can express the observed signal intensities as:

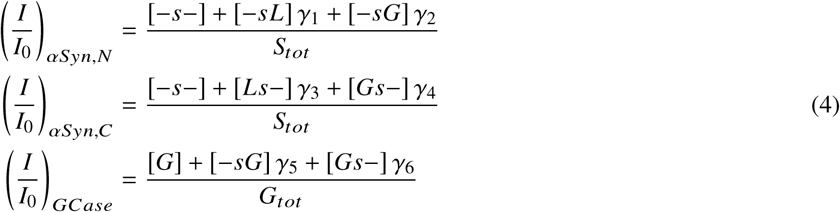

The most complex model described by this picture is parametrized by 19 values (9 equilibrium constants including the ternary species and 10 attenuation constants). This model was found to provide the best fit to the data. But as we have 21 individual signal intensity measurements, to apply this model carries a strong risk of over-fitting. We tested a number of models both including and excluding ternary complexes, and with varying numbers of species included that are proposed to be NMR observable (non zero *γ* factors).

Following the principles of sensible fitting, we sought to determine the minimally complex model where the reduction in *χ^2^* to more complex models is determined to be statistically insignificant by application of an F-test. The *χ^2^* was defined as:

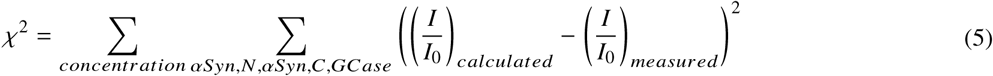

Such testing revealed that no significant improvement was obtained if we incorporate 1) lipids binding the C-terminus of *α*Syn, 2) GCase binding the N-terminus of GCase, nor 3) any of the ternary complexes. Moreover, there was no statistical benefit to including observation of GCase signal when it is bound to lipids. This results in the following simplified model that we conclude gives the best statistical fit to the data, with three equilibrium constants:

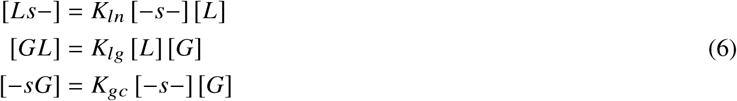

And three attenuation exchange factors whose origin comes from chemical exchange (*γ*):

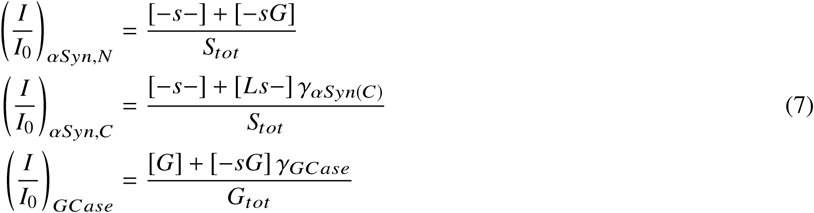

This model (Fig S6, model 3, S7) is not significantly improved over a model where we set *γ*3 to 1, which amounts to the N-terminus of *α*Syn contributing to the observed signal intensity equally to free *α*Syn. Overall this model (Fig S6, model 4) is parametrized by 5 constants and predicts the existence of 3 binary complexes in addition to the 3 free species. This model gives an excellent description of all signal intensities recorded over the wide range of combinations of concentrations tested. Incorporating additional parameters did not result in a statistically significant improvement in the fitting. The formulation of the signal intensities successfully account for the observed increase in diffusion coefficients (Fig S4) as it predicts bound species contribute the spectrum albeit with reduced signal intensity when compared to the free species.

The code to perform this analysis was written in python, where it was determined that a stimulated annealing approach was necessary to find the global optimum. Uncertainties in fitting parameters were derived from a Monte-Carlo method, with between 20% and 30% uncertainty in each parameter. The uncertainty is relatively high. The trends in these parameters however span orders of magnitude (Fig 6B) and so we obtain meaningful mechanistic insight with this approach. The high uncertainties also imply that incorporation of additional information into the model, for example a more detailed description of chemical exchange parametrized with additional data could further refine the model. The optimal model does not include ternary complexes. We note that this does not mean ternary complexes cannot form, just that we see no statistically significant reason in the data to justify their inclusion to quantitatively explain the data, suggesting if they are formed they are highly unstable. The formation of ternary complexes has been previously inferred from fluorescence changes (59) though this finding is not supported by our data on the mixtures analysed here.

## RESULTS

### GCase inhibits *α*Syn amyloidogenesis under lysosomal conditions

To mimic the amyloidogenesis of *α*Syn in the lysosome, *α*Syn aggregation was monitored using the amyloid specific dye thioflavin-T (ThT) at pH 5 in the presence and absence of both GCase and assemblies of anionic lipids (Figure 1) (44, 50). The aggregation of *α*Syn at pH 5 was markedly faster in the presence of DMPS, reducing the lag time from 3 days (red), to 1 (blue) (Figure 1B/C). Although the aggregates bound ThT and possessed fibrillar morphological aspects, discrete fibrils were not observed in electron micrographs (Figure 1B). Remarkably, adding GCase effectively prevented aggregation both in the presence (cyan) and absence (green) of DMPS, demonstrating that it is a potent inhibitor of *α*Syn amyloidogenesis under lysosomal conditions.

### GCase and anionic lipids compete for opposite termini of monomeric *α*Syn

In order to understand the molecular basis of this interaction, we recorded fingerprint ^15^N-^1^H HSQC NMR spectra in absence and presence of DMPS, POPG, SDS, DDM, and tween under otherwise identical lysosomal conditions. In a ^1^H-^15^N HSQC spectrum, one resonance is observed for each distinct HN environment in the molecule. ^1^H NMR resonances fell predominantly in the region 8-8.5 ppm, confirming the expected disordered character of *α*Syn (Fig 2A).

**Figure 2:**
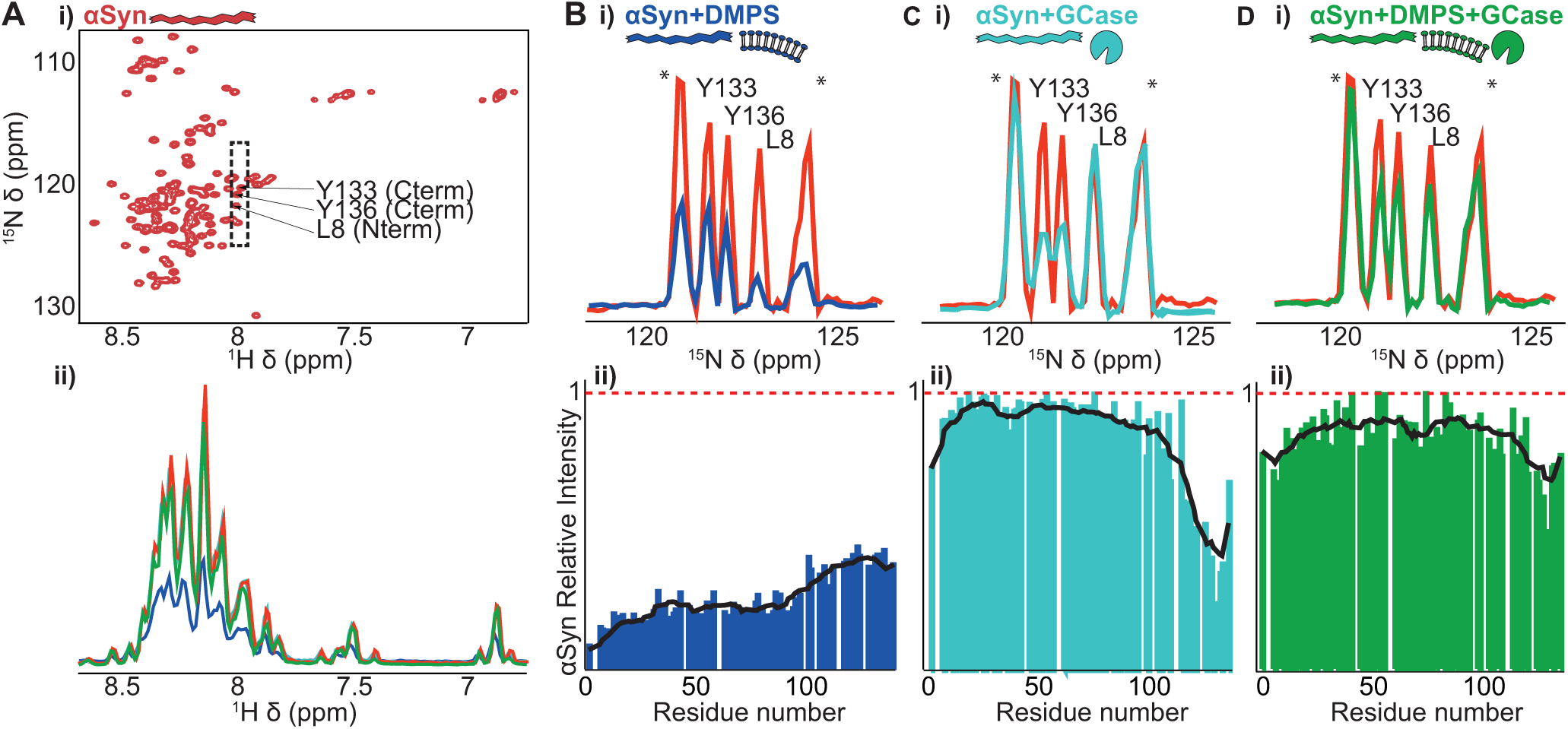
GCase rescues *α*Syn from lipid binding under lysosomal conditions. **A**) i) A lack of *α*Syn signal dispersion in a 2D ^1^H-^15^N HSQC spectrum confirms its intrinsically disordered character. ii) Incubating *α*Syn (red) with the anionic lipid DMPS (blue) attenuates the intensity of all signals due to formation of high molecular weight complexes, whereas the addition of GCase (green) recovers it. In **B**, **C**, and **D**, the (i) same 2D ^1^H-^15^N HSQC region (see box in Ai) and (ii) corresponding *α*Syn signal intensities are shown in the absence or presence of DMPS or GCase, respectively, relative to the signal intensity of free *α*Syn. Addition of GCase significantly increases the amount of free *α*Syn present in solution.

In the presence of the anionic taurochlorate, DMPS and 1:1 mixtures of DDM with SDS and POPG (Fig S1) a residue specific reduction in signal intensity was observed, with no significant change in either resonance frequencies or observed linewidths (fig 2Aii, S2, S3). In line with previous observations, N-terminal *α*Syn residues showed a greater decrease in signal intensity than those in the C-terminus (Fig 2B, S2). Moreover, a higher lipid ratio decreased the N-terminal NMR signal intensities further (Fig S2). By contrast, micelles formed from the non-ionic detergents DDM and tween resulted in no change in signal intensity (Fig S3). In the case of the anionic lipids, the observed diffusion coefficients of *α*Syn were significantly reduced (Fig S4). Together, the loss in signal intensity can be explained by the formation of *α*Syn lipid complexes, where any structured portion of the molecule tumbles too slowly to be observed directly by solution state NMR.

Next we determined the effect of GCase on *α*Syn in a 1:1 mixture of both components. Contrasting with lipid binding, a signal intensity reduction was observed localized to the C-terminus of *α*Syn (Fig 2C). Incubation of *α*Syn in the presence of lipids and GCase (figure 2D) resulted in a recovery of signal intensity in both the N- and C-termini. Furthermore, NMR diffusion coefficients of *α*Syn returned to values similar to those measured for the isolated form (Fig S4). These results indicate that GCase substantially rescues *α*Syn from its lipid-bound amyloidogenic state.

To allow for a quantitative comparison of the effects at the N- and C-termini of *α*Syn, the signal intensities of the first and last 20 residues were integrated for each of the mixtures of *α*Syn with GCase and lipid SUVs/micelles (Fig 3,S2C,S3F/G). Although the magnitudes of the interaction of *α*Syn with different lipids vary, GCase was always found to rescue the signal intensity of *α*Syn (Fig 3).

**Figure 3:**
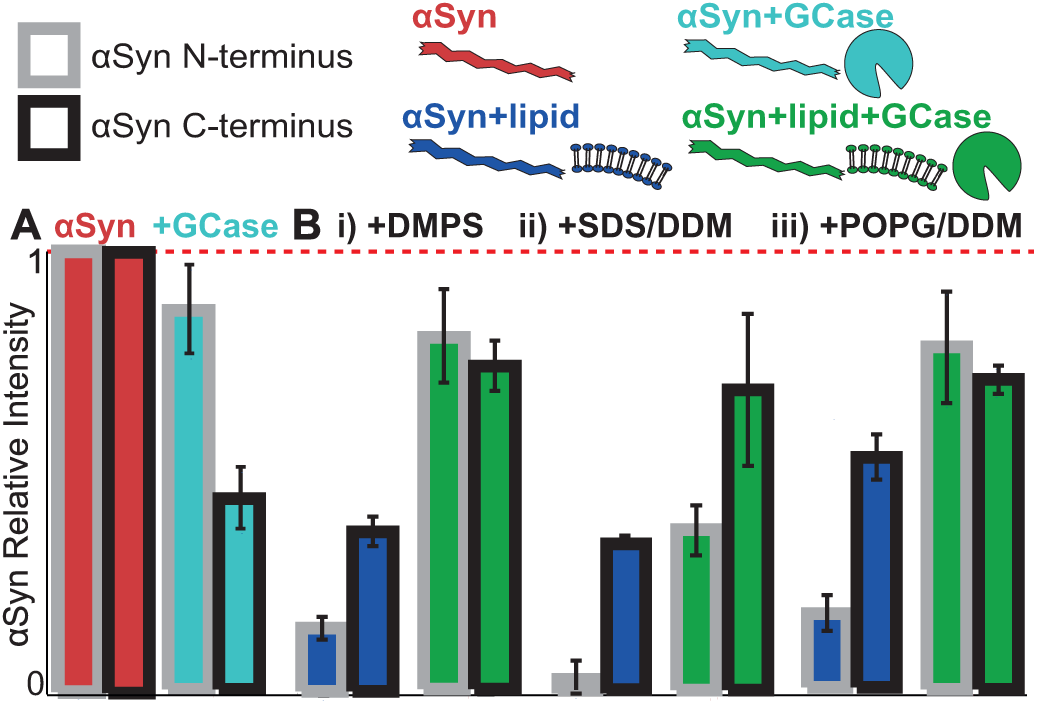
Integrated relative signal intensities and between residue standard deviations of the 20 most N- (grey) and C- (black) terminal *α*Syn residues in the presence of GCase (cyan), anionic lipid SUVs/micelles formed from DMPS,SDS/DDM, or POPG/DDM (blue) and their combination (green). In each case there is substantial recovery in the observed signal intensity by adding GCase (green).

### GCase destabilises mature *α*Syn amyloid fibrils

We sought to examine the effect of GCase on pre-formed amyloid fibrils. At elevated temperatures and concentrations (200 *μ*M, 45°C) ^15^N labelled *α*Syn showed increased ThT fluorescence (Fig 4A), and discrete amyloid fibrils were observed by EM (Fig 4B) (44). A ^1^H-^15^N HSQC NMR spectrum of these samples at pH 5 revealed a small population of flexible C-terminal residues observable by NMR, a finding in agreement with proteolysis and solid state NMR studies (35, 61, 62). Most remarkably, addition of GCase to these samples recovered signal from N-terminal residues (Fig 4C), indicating a destabilization of the amyloid form and partial rescue of monomeric *α*Syn by GCase.

**Figure 4:**
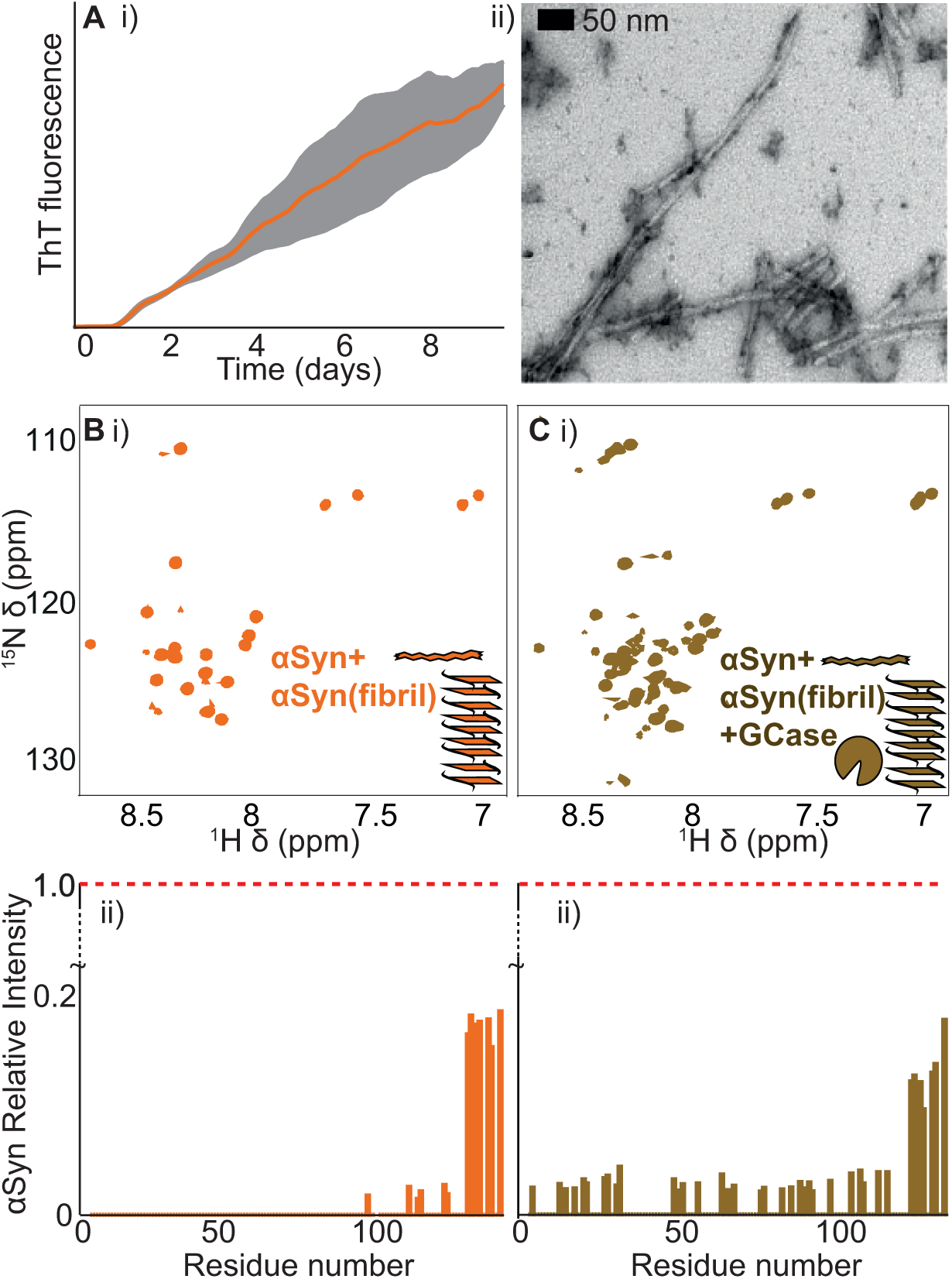
**A**) i) The *α*Syn aggregation kinetics were monitored by a thioflavin-T fluorescence assay at elevated temperature and concentration (45°C, 200 *μ*M) at pH 5. ii) These conditions result in discrete amyloid fibrils, as observed by EM. **B/C**) Amyloid fibrils were purified by ultra centrifugation, and its ^1^H-^15^N HSQC NMR spectrum was taken (Bi, orange). GCase was then added and the spectrum was obtained (Ci, brown). All spectra were acquired under lysosomal conditions (pH 5). ii) The relative intensity of the oserved resonances as a function of residue position in the *α*Syn sequence. Incubation with GCase results in a significant release of free *α*Syn.

### GCase interacts with *α*Syn near its active site

Post-translational modifications are vital for active and stable GCase (30, 63, 64). While protein with these modifications can be prepared from human cell lines, introducing the necessary isotopes for NMR analysis during expression is prohibitively expensive. Moreover, the high molecular weight of GCase (60 KDa) would significantly broaden the width of observed NMR resonances in a protonated sample. To overcome this, we introduced ^13^ C ^1^ H _3_ methyl groups to specific locations on the protein surface. The large number of equivalent protons and rapid internal dynamics result in to sharp resonances in NMR spectra, making such probes highly desirable for large molecular weight proteins.

To incorporate ^13^C^1^H_3_ methyl groups into human cell derived GCase containing human post translational modifications, we employed chemical modification. Amine methylation is a routine strategy in crystallography and although it leads to covalent modification, protein structure and interactions are not generally perturbed (49, 54, 65–69). To maximize coverage of the protein surface, we targeted lysine and arginine residues using ^13^C labeled formaldehyde (67, 70) rather than targeting the smaller number of cysteine residues (71).

GCase (Shire PLC) was produced from human cell lines and then incubated with ^13^C formaldehyde to produce lys^13^C^1^H_3_ and arg^13^C^1^H_3_ enriched GCase (methods). The successful modification of specific lys and arg residues was confirmed by liquid chromatography tandem mass spectrometry (LCMSMS) (Fig S5) with no significant change in activity (Fig S5D). Most resonances in the ^1^H-^13^C HMQC spectra were heavily overlapped due to a combination of low chemical shift dispersion, a large number of labels and the intrinsically slow tumbling of the 60 kDa protein leading to broadened resonances (Fig 5B). By analysis of additional 2D ^1^H-^13^C HMQC spectra and comparison to LCMSMS data under different methylation conditions, however, one resonance could be unambiguously assigned to the active side residue R120 (72) (Fig S5).

**Figure 5:**
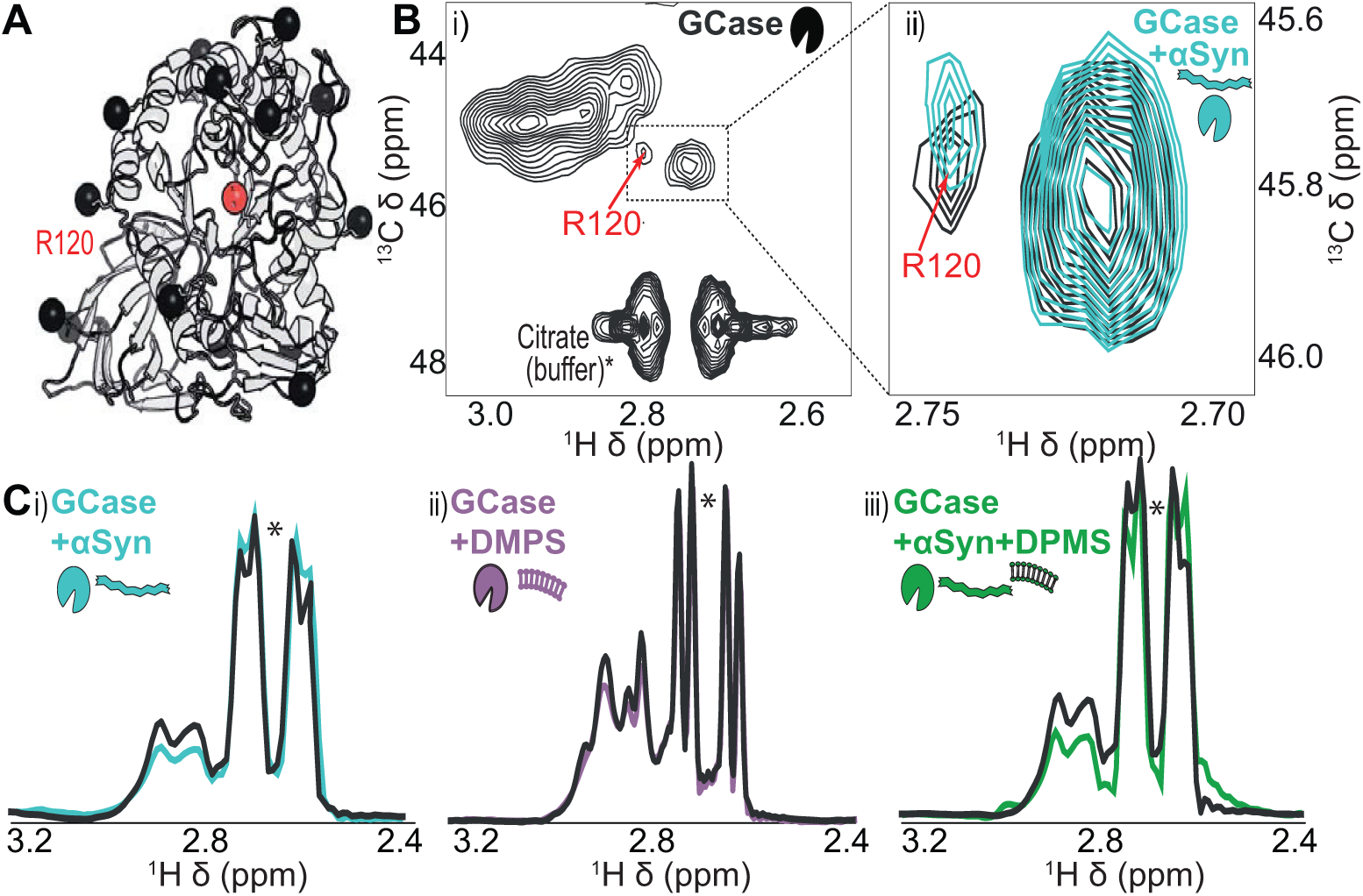
**A**) Lysine and arginine residues on the surface of GCase (PDB 1OGS) were modified by incubation with ^13^C labelled formaldehyde (black spheres, see Fig S5). **B**) i) 2D ^1^H-^13^C HMQC NMR spectrum of the labelled residues. Natural abundance of ^13^C citrate results in sharp resonances in the indicated positions (*). While the majority of resonances overlap, one distinct peak from R120 could be distinguished and unambiguously assigned using LCMSMS (Fig S5). ii) After incubation with *α*Syn, the locations of all the resonances were largely unchanged. Only the resonance corresponding to R120 was found to move, suggesting a direct interaction between this site and *α*Syn. **C**) ^1^H projections of 2D spectra after arginine methylation (black) and mixtures with *α*Syn and anionic lipids (cyan/purple/green). As GCase samples were prepared independently, small fluctuations in spectra between samples were observed (black i,ii,iii). The small differences in signal intensity from the citrate in projection are due to truncation artefacts in the spectra, and do not reflect changes in citrate concentration. The uniform loss of signal intensity from resonances originating from GCase after incubation with *α*Syn, DMPS or a mixture of both indicates the formation of high molecular weight species that are not directly observed.

The ^13^C^1^H_3_ methyl groups were used to monitor the effects of both lipids and *α*Syn on GCase (Fig 5C,6A). The addition of either *α*Syn or DMPS resulted in decreased GCase signal intensity. Most notably, a modest yet significant change in the chemical shift of the peak arising from R120 was observed on mixing with *α*Syn, whereas the resonance frequencies of the remaining residues were largely unchanged. This suggests that the interaction site is in the vicinity of this location. As R120 is close to the active site, this observation is consistent with *α*Syn reducing the activity of GCase (20,59) and with FRET data (59).

**Figure 6:**
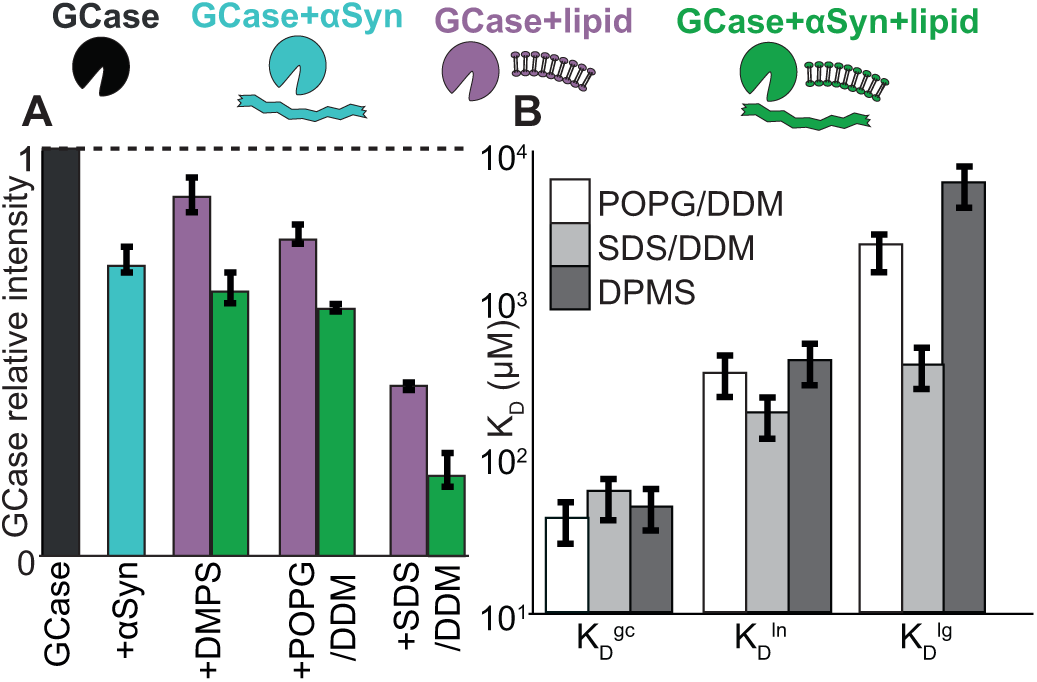
**A**) Summary of the changes in resonance intensity of ^13^C^1^H_3_ labelled GCase on incubation with *α*Syn (cyan), lipids DMPS, POPG/DDM, SDS/DDM (purple), and both *α*Syn and lipid (green). **B**) Quantitative *K_D_* values for the three processes from fitting the combined *α*Syn, GCase and lipid binding data to optimal model (Fig S6,S7). The dissociation constant for GCase/*α*Syn complex formation was independent of lipid choice. The specific *K_D_* s for both *α*Syn and GCase lipid binding are highly dependent on the choice of lipid, and are highly correlated. The strongest binding was observed with SDS/DDM, and the weakest with DMPS.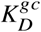
describes the interaction between the C-terminus of *α*Syn and GCase,
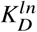
describes the interaction between the N-terminus of *α*Syn and lipids, and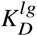
describes the interaction between GCase and lipids.

### GCase prevents *α*Syn aggregation through a competitive equilibrium

Our data allows us to follow the interactions between *α*Syn, lipids and GCase from the perspective of *α*Syn and GCase. The loss of signal intensities in both *α*Syn and GCase were both reversible, and dependent on the concentrations of the three species (Fig 2,3,4,6,S2,S3) revealing that these two species on the timescale of the NMR experiments are in competition with lipids. We sought to determine whether a simple equilibrium model could explain our combined datasets. To do these, we needed to analyse the possible origin of intensity losses in our NMR spectra. Having explored a large range of different concentrations of the three reagents, coupled to the large number of probes reporting on various structural regions of both *α*Syn and GCase, our datasets are well suited to providing significant insight into the mechanism by which the three interact.

As their slow tumbling prohibits a direct observation of SUVs and GCase in ^1^H NMR spectra, we would also not expect to observe regions of *α*Syn directly bound to either lipids or GCase. Consequently, complex formation is registered in the form of a loss of NMR signal intensity. If only one terminus of *α*Syn bound SUV or GCase, the other terminus would remain flexible and detectable by NMR. This allows not only for a differential analysis of both termini but also a quantification of attenuated NMR signal intensities.

As we did not observe any line broadening, chemical shift perturbations(32, 33, 73), or variations in CPMG-derived transverse relaxation rates (data not shown) of *α*Syn residues, our results suggest that chemical exchange is in slow or slow-intermediate limits (32, 73, 74). We account for these effects in the NMR spectra by allowing regions of *α*Syn that are expected to retain significant flexibility to contribute to the NMR spectrum in a manner that is proportional to the concentration of complex formed, thought reduced from that of the free state (methods). This exchange regime is also consistent with the reduced diffusion coefficients observed upon binding (Fig S4).

Subsequently, we tested a wide range of possible binding models (Fig S6) and used F-tests to identify the simplest model explaining all of the data, while avoiding over-fitting (methods). The simplest model tested considers only binary complexes and does not distinguish between binding to the N- and the C-termini of *α*Syn, whereas the most complex model included all possible ternary complexes (Fig S6). The optimal model explains all of our data quantitatively (Fig. S7), considers all interactions with DMPS, POPG/DDM, and SDS/DDM, is consistent with the decreased diffusion coefficients (Fig. S4). It allows for both GCase and the N-terminus of *α*Syn to bind lipids, and the C-terminus of *α*Syn to bind GCase. It is parametrized by three dissociation constants for the formation of complexes between GCase and lipids 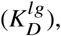 the C-terminus of *α*Syn and GCase 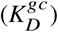 and the N-terminus of *α*Syn and lipids 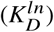 (Fig S6,S7). It also takes attenuated NMR signal intensities of an *α*Syn terminus into account when the opposite terminus is involved in complex formation (methods).

Overall, the model reveals simple equilibrium binding where the N- and C-termini of *α*Syn are in competition with lipids and GCase, respectively (Fig 6B,S7). The
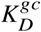
for the GCase/*α*Syn(C) interaction was ca. 60 *μ*M for all lipids tested, whereas the
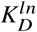
for the *α*Syn(N)/lipid interaction followed the degree of charge on the lipid head group in the sequence DMPS, POPG/DDM, SDS/DDM. It is interesting to note that the lipids that bound GCase more tightly also had a higher affinity for *α*Syn suggesting that *α*Syn and GCase compete for similar binding sites on the surface of lipids.

All of the data can be quantitatively summarized on a free energy surface (Fig 7). Free *α*Syn (red) will form amyloids (orange) if left for sufficient time. This process is greatly accelerated by anionic lipids (blue, Fig 1). GCase effectively competes with lipids for binding of *α*Syn (cyan). By reducing the population of lipid bound complexes GCase prevents the build-up of amyloidogenic precursors to inhibit amyloid formation. If left for long enough, stable amyloid formation is expected to become inevitable (75). Yet the formation of rapid and relatively weak contacts between GCase and *α*Syn greatly curtails amyloidogenesis.

**Figure 7:**
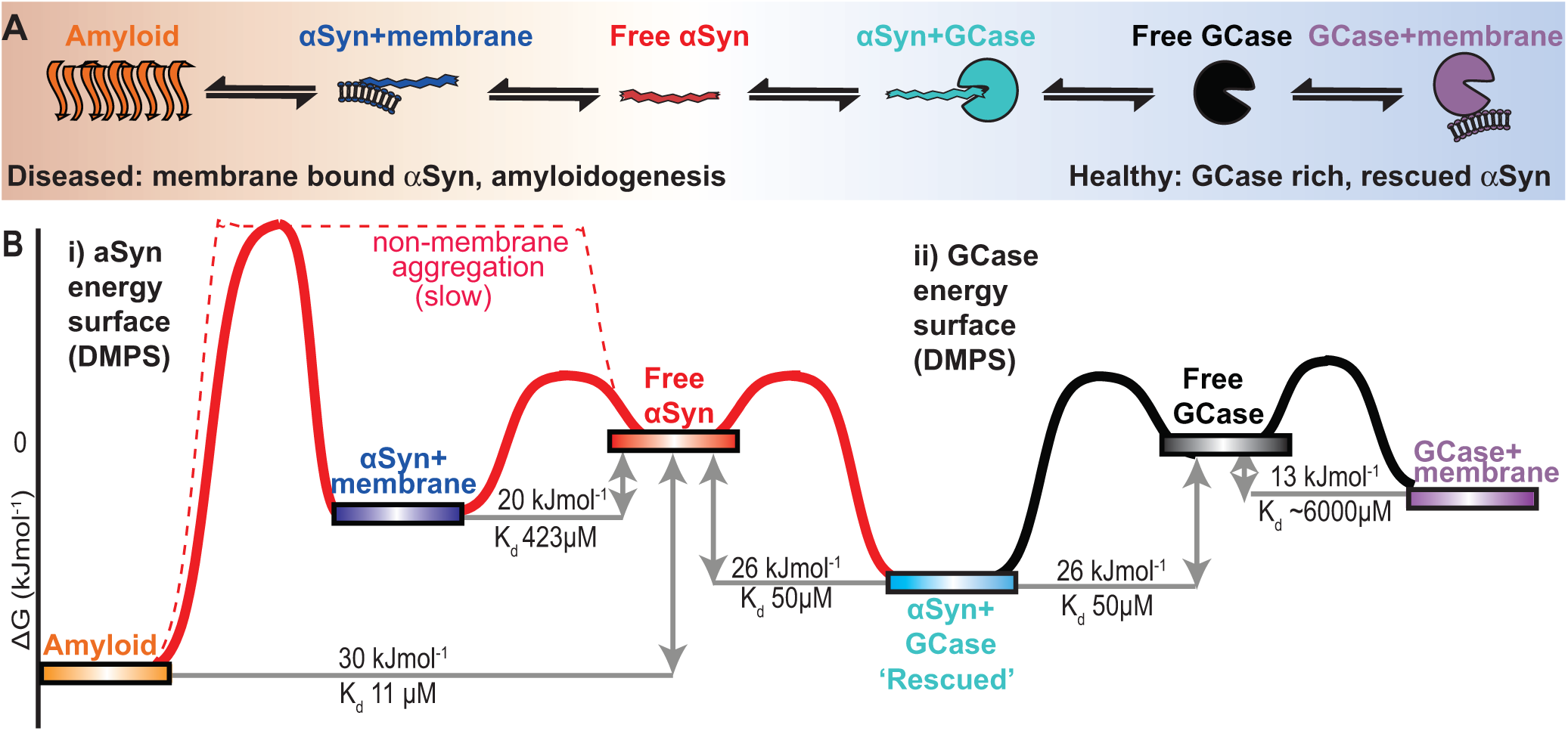
A quantitative equilibrium competition model between *α*Syn, GCase and lipid well explains all of our data. *α*Syn aggregation is accelerated by the presence of anionic lipids (Fig 1). GCase binding greatly inhibits this process, inhibiting *α*Syn amyloidogenesis (Fig 1). **B**) A schematic free energy surface for DMPS displays how all complexes are related. The presence of *α*Syn-GCase complexes effectively rescues *α*Syn by preventing their build up on the surface of membranes. The energy landscape is shown from the perspective of *α*Syn (i) and from GCase (ii). All binding reactions came to equilibrium at maximum on the timescale of a few minutes, whereas *α*Syn aggregation occurs on a timescale of days under these conditions. The uncertainties of the specific free energies can be propagated from the values in figure S7A.

## DISCUSSION

Here, we demonstrate that GCase directly inhibits amyloidogenesis of *α*Syn both in the presence and absence of anionic lipids under lysosomal solution conditions (Fig 1). Quantitative analysis of the changes in both GCase and *α*Syn reveals a molecular ‘tug of war’ mechanism in which GCase and anionic membrane mimics are in competition for the C- and N-termini of *α*Syn respectively (Fig 2). The strongest interaction identified here between GCase and the C-terminus of *α*Syn inhibits amyloidogenesis, revealing a protective role for GCase in PD pathology. Though not explicitly characterised here, complexes between *α*Syn and GCase clearly inhibit the formation of *α*Syn aggregation nuclei both on and off lipids (Fig 1). These results demonstrate that GCase can function as a molecular chaperone, reversibly binding *α*Syn and both inhibiting the formation and reducing the stability of amyloid formed from *α*Syn.

Central to the determination of this mechanism was the need to study human cell derived GCase complete with relevant post-translational modifications. The methylation methodology we employ is generally applicable for studies of biologically relevant proteins without the need to be prepared in lower organisms.

In a healthy lysosome, a GCase rich lysosome is able to degrade both monomeric *α*Syn and *α*Syn oligomers (6, 15, 17). In PD, depletion of GCase in the lysosome is associated with the build-up of amyloid and enhanced susceptibility to Parksinson’s disease (5, 8, 11, 14, 15, 17, 20, 21, 76, 77). *α*Syn amyloidogenesis in the lysosome is enhanced by both GCase depletion, and the presence of anionic lipids (4, 5, 14, 15, 17, 20, 27). GCase depletion also enhances the transfer of *α*Syn oligomers between cells by the lysosomal system (20, 25, 26). The presence of anionic lipids and mutations that deplete the lysosomal GCase concentration are both significant risk factors in PD. Our model explains these observations, predicting a shift in equilibrium favouring increased lipid-induced amyloid formation in the case of GCase depletion under lysosomal conditions. Furthermore, GCase’s destabilising role on amyloid fibrils explains why the depletion of GCase results in an enhanced transfer of *α*Syn oligomers between lysosomes of adjacent cells (20, 25, 26).

Taken together, our quantitative analysis explains the protective effect of GCase on PD models and why its genetic and sporadic depletion results in an increased risk of PD (13, 78). We demonstrate that wild-type GCase can function as a molecular chaperone, inhibiting *α*Syn amyloid formation under lysosomal conditions. These results provide a molecular framework to interpret the potential therapeutic benefits of GCase in Parkinson’s disease.

## CONCLUSION

The lipid-accelerated deposition of *α*Syn into amyloid rich Lewy bodies is a key hallmark of Parkinson’s disease. Recently, lysosomal enzyme glucocerebrosidase has been implicated in sporadic and non-sporadic Parkinson’s disease. Here, we have shown by solution state NMR the protective role of human-cell derived GCase in inhibiting *α*Syn amyloid formation and disrupting pre-formed amyloid fibrils under lysosomal conditions. We have further localised interaction sites using human cell derived GCase. And proposed a “tug of war” mechanism where GCase and lipids compete for different termini of *α*Syn, explaining the relationship between GD and PD.

## AUTHOR CONTRIBUTIONS

MB led the design and execution of the research with AB, AB led the statistical modelling of the NMR data with MB. HM provided expertise in electron microscopy and RG aided with molecular biology techniques.

## ACKNOWLEDGEMENTS

AJB thanks the BBSRC for a David Phillip’s Fellowship. MB thanks the BBSRC for Doctoral Training Partnership funding. The authors thank T.R. Alderson for the *α*Syn plasmid and Shire PLC for GCase (VPRIV) samples. The Division of Structural Biology is a part of the Wellcome Trust Centre for Human Genetics, Wellcome Trust Core Grant Number 090532/Z/09/Z.

